# Functional insights into the effect of feralisation on the gut microbiota of cats worldwide

**DOI:** 10.1101/2024.09.04.611329

**Authors:** Ostaizka Aizpurua, Amanda Bolt Botnen, Raphael Eisenhofer, Iñaki Odriozola, Luisa Santos-Bay, Mads Bjørn Bjørnsen, MTP Gilbert, Antton Alberdi

## Abstract

Feralisation, the process by which domesticated organisms revert to a wild state, is a widespread phenomenon across various species. Successfully adapting to a new environment with different access to food, shelter, and other resources requires rapid physiological and behavioural changes, which could potentially be facilitated by gut microbiota plasticity. To investigate whether alterations in gut microbiota support this transition to a feral lifestyle, we analysed the gut microbiomes of domestic and feral cats from six geographically diverse locations using genome-resolved metagenomics. By reconstructing 229 draft genomes from 92 cats, we identified a typical carnivore microbiome structure, with notable diversity and taxonomic differences across regions. While overall diversity metrics did not differ significantly between domestic and feral cats, hierarchical modelling of species communities, accounting for geographic and sex covariates, revealed distinct taxonomic and functional profiles between the two groups. While taxonomic enrichment was balanced, microbial functional capacities were significantly enriched in feral cats. These functional enhancements, particularly in amino acid and lipid degradation, correspond to feral cats’ dietary reliance on crude protein and fat. Additionally, functional differences were consistent with behavioural contrasts, such as the more aggressive and elusive behaviour measured in feral cats compared to the docile behaviour of domestic cats. Finally, the observed enrichment in short-chain fatty acid, neurotransmitter, and vitamin B12 production in feral cats aligns with improved cognitive function and potentially contributes to their heightened aggression and elusiveness. Our findings suggest that microbiome shifts may play a significant role in the development of physiological and behavioural traits advantageous for a feral lifestyle, supporting the adaptive success of feral cats in the wild.

## Introduction

Feralisation is the process by which a once-domesticated organism detaches from the anthropic environment (Henriksen et al., 2018). While possessing some distinct characteristics, feralisation can generally be perceived as the counterpoint to domestication (Price, 1984). Many species initially domesticated by humans have subsequently given rise to feral populations (Gering et al., 2019), often leading to adverse impacts on both human settlements and biodiversity (Bonacic et al., 2019; Medina et al., 2011; Palmas et al., 2017). Despite the profound implications of this phenomenon, feralisation remains a relatively understudied process compared to its counterpart, domestication.

One of the animal species that is commonly found in feral form is the house cat (*Felis silvestris catus*). The domestication of cats is thought to have taken place in the Near East at the onset of the Neolithic period (Driscoll et al., 2007). This likely occurred concurrently with the rise of agriculture, as cats became valuable assets in controlling rodent populations around grain stores (Clutton-Brock, 1990; Gross, 2020). Since then, cats have spread alongside human populations across most of the planet (Baca et al., 2018; Ottoni and Van Neer, 2020). In every region they have colonised, cats have demonstrated an exceptional ability to thrive outside of domestic settings, forming countless feral cat populations worldwide (Bradshaw et al., 1999). Therefore, while domestication happened in a specific geographic area and time point, feralisation has happened and is continuously happening all over the world. This is partly due to the fact that modern non-pedigree domestic cats still closely resemble their wild ancestors both genetically and morphologically, retaining a significant portion of their wild behavioural repertoire, including proficient hunting skills (Bradshaw, 2006; Cecchetti et al., 2021). Additionally, feral cats typically display behaviours more akin to their wild relatives, such as an aversion to humans (Levy et al., 2003). The feralisation of cats poses a significant threat to the environment, as feral cats have been implicated as the primary cause of at least 14% of global bird, mammal, and reptile extinctions (Dueñas et al., 2021; Medina et al., 2011).

Feralisation success is often linked to phenotypic plasticity (Gering et al., 2019), as feral animals must adapt to significantly more variable and unpredictable environments compared to their domestic counterparts. In this regard, there is growing interest in the potential role that the gut microbiota may play in assisting such transitions. In general, exposure to new environments can lead to the acquisition of novel intestinal microorganisms, and these can interact with the host in diverse ways (Candela et al., 2012). While traditional microbiological and veterinary research has largely concentrated on pathogens, there is an increasing recognition of the beneficial effects of gut microorganisms (Lee and Hase, 2014; Ma et al., 2023). Researchers have hypothesised that these microorganisms can influence the phenotypic plasticity of animals, thereby enhancing their ability to thrive in diverse environments (Alberdi et al., 2016; Henry et al., 2021; Moeller and Sanders, 2020; Rosenberg and Zilber-Rosenberg, 2022; Voolstra and Ziegler, 2020). For example, variability in gut microbiota is acknowledged as one of the most effective mechanisms for adapting to dietary changes (Teullet et al., 2023). Moreover, changes in gut microbiota have been linked to animal behaviour, as gut microorganisms are known to produce and metabolise neurotransmitters and hormones that can influence neurological processes (Forsythe et al., 2014; Johnson and Foster, 2018; Mayer, 2011). In fact, studies on domestic chickens and foxes kept in controlled conditions have demonstrated how certain microbiome features are associated with different behavioural responses to humans (Puetz et al., 2024, 2021). These observations raise the hypothesis that the differentiation in behaviour often observed when comparing feral and domestic individuals, may be partially influenced by differences in their gut microbial communities.

In this study, we elected to explore how the gut microbiota of cats changes in association with feralisation. Specifically, to explore their potential implications in facilitating the process and shaping behavioural changes in hosts, we used genome-resolved metagenomics to compare the taxonomic and functional features of gut microbiomes of 92 domestic and feral cats, sampled in six different locations across the world. With this data, we tested whether feral cats derived from domestic populations recovered an extended repertoire of microbial functions compared to domestic cats in the same region, and whether these changes were in line with the observed behavioural phenotypes.

## Results

### A global catalogue of cat-associated microbial genomes

We generated 298.1 GB of shotgun sequencing data (3.3±1.4 GB/sample) from faecal samples of 96 cats collected across six countries (Figure 1a). Metagenomic assembly and binning of this dataset yielded a total of 229 non-redundant metagenome-assembled genomes (MAGs) (Figure 1b). The average CheckM completeness of these MAGs was 90.02 ± 7.79%, with an average CheckM contamination of 1.32 ± 1.18% (Supplementary Fig. S1).

**Figure 1.**
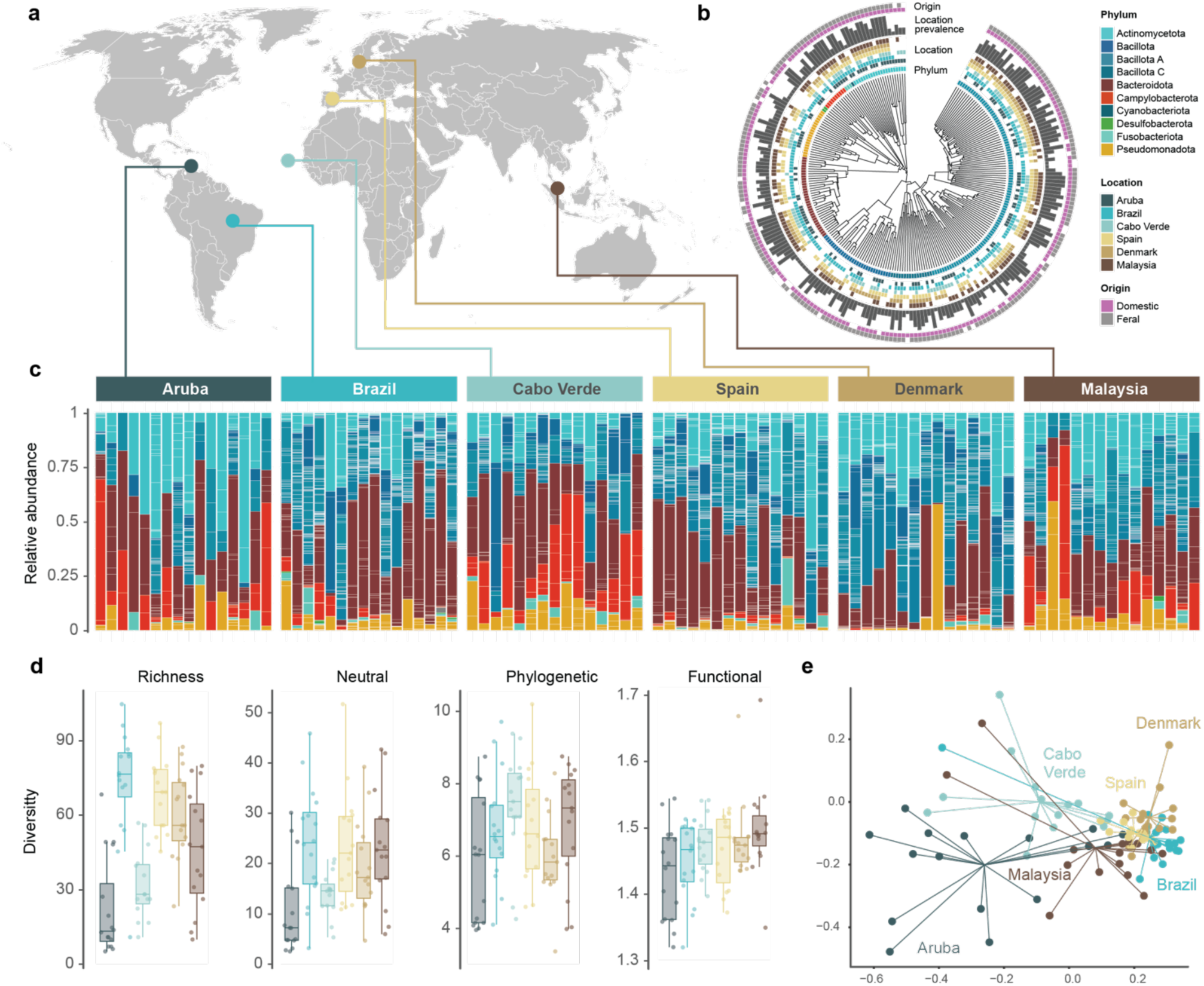
Overview of the geographic locations and genome catalogue characteristics. **a)** World map with the geographic locations of the samples. **b)** Reference metagenome-assembled genome (MAG) catalogue generated from the cat faecal samples, with outer rings indicating detection of MAGs in different geographic regions and origin (domestic or feral). **c)** Taxonomic composition of the faecal samples, each tile representing a MAG coloured by taxonomic phyla. **d)** Alpha diversity differences across geographic regions, based on four different Hill numbers diversity metrics. **e)** Ordination of richness-based compositional differences across samples, coloured and grouped by location.

The MAGs were taxonomically assigned to 10 bacterial phyla (Figure 1b), with nearly half of them belonging to Bacillota A. Despite their phylogenetic relatedness, MAGs belonging to Bacillota A displayed extensive functional variability, covering over half of the functional space in the functional ordination (Supplementary Fig. S2). Bacterial communities were dominated by Bacteroidota (30.62±17.35%), Bacillota_A (21.47±10.53%), and Actinomycetota (17.85±14.45%), with the remaining bacterial phyla with average value with less than 9% (Figure 1c). At the family level, Bacteroidaceae (28.71±16.84%), Lachnospiraceae (12.18±8.03%), Coriobacteriaceae (9.96±9.09%), Helicobacteraceae (4.59±7.93%), and Megasphaeraceae (4.37±7.38%) were the five most abundant in the gut microbiome of cats (Supplementary Fig. S3). Furthermore, *Prevotella* (13.11±13.07%), *Collinsella* (9.96±9.08%), *Phoecaeicola* (8.98±8.41%), *Bacteroides* (4.85±4.99%) and *Megasphera* (4.33±7.35%) the five most abundant bacterial genera (Supplementary Fig. S4).

### Differences across localities

While 66 MAGs (29%) were present across all locations, the rest were specific to one or a few locations (Supplementary Fig. S5). Brazilian samples exhibited the highest number of MAGs, including 20 unique MAGs, with Cabo Verde showing both the fewest total and unique number of MAGs. The spread of bacteria was not affected by the lifestyle of the cats (MANN-WHITNEY; W = 25706, p-value = 0.713), with MAGs from domestic and feral cats being present in 3.12±1.93 and 3.19±2.03 locations, respectively. Diversity metrics also varied across geographic locations, with Aruba and Cabo Verde exhibiting the lowest neutral diversities, yet similar phylogenetic and functional metrics as the rest of the locations (Figure 1d).

The location of the cats explained most of the compositional variability of the gut microbiota (Figure 1e). Over 20% of the variability of the neutral (PERMANOVA; R^2^=0.221, p-value=0.001) and phylogenetic (PERMANOVA; R^2^=0.268, p-value=0.001) diversity was explained by the location and 58% of the functional variability (PERMANOVA; R^2^=0.584, p-value=0.001). Spain and Denmark exhibit more similar microbial communities than the rest of the localities (pairwise-Permanova: R^2^=0.0972, p-adjusted=0.075). Despite geographical differences, cats from Brazil had more similar gut microbiota to cats from Spain (pairwise-Permanova: R^2^=0.0630, p-adjusted=0.555) than from the nearby Aruba (pairwise-Permanova: R^2^=0.1377, p-adjusted=0.015). Cats from Aruba, Cabo Verde and Malaysia exhibited a more distinct microbial community than those from Denmark, Brazil, and Spain, which clustered closely together (Supplementary Fig. S6).

### Functional microbiome differences between domestic and feral cats

Prior to comparing the microbiomes of the feral vs domestic cats, we validated that cat origin did correlate with behavioral traits. Specifically, feral cats exhibited less acceptance (Χ^2^ = 24.525, df = 1, *p*-value = 7.337e-07) and more hisses (Χ^2^ = 11.221, df = 1, *p*-value = 0.0008), bites (Χ^2^ = 11.808, df = 1, *p*-value = 0.0006) and retreats (Χ^2^ = 5.4412, df = 1, *p*-value = 0.0197) towards humans compared to domestic cats. When comparing the gut microbiota of these behaviourally different groups we observed that overall alpha (Mixed models, p>0.05) and beta diversity (PERMANOVA, p>0.05) analyses did not yield any significant differences in microbiome diversity and composition between feral and domestic cats (Figure 2a-b). Accordingly, the only bacteria unique to either of the two origins were rare species found in just one location, with no widespread MAGs specific to either domestic or feral origins. However, when considering relative abundances, hierarchical modelling of species communities (HMSC) revealed that 61 MAGs were associated with domestic cats while 34 MAGs were associated with feral cats (Figure 2c-d). MAGs belonging to a range of bacterial phyla were enriched in domestic cats, including Pseudomonadota (11 MAGs from CAG-239, Burkholderiaceae, and Succinivibrionaceae), Campylobacterota (3 MAGs from Helicobacteraceae), Bacteroidota (9 MAGs from Bacteroidaceae), Actinomycetota (18 MAGs from Actinomycetaceae, Atopobiaceae, Bifidobacteriaceae, Coriobacteriaceae, Mycobacteriaceae), Bacillota A (11 MAGs from Butyricicoccaceae, Oscillospiraceae, Peptoniphilaceae and Ruminococcaceae) and Bacillota C (8 MAGs from Acidaminococcaceae, Dialisteraceae, Megasphaeraceae), Desulfobacterota (1 MAG from Desulfovibrionaceae). In feral cats, MAGs belong to Bacillota A (14 MAGs from Acutalibacteraceae, Anaerotignaceae, Clostridiaceae, Lachnospiraceae, Peptostreptococcaceae, and UBA1381), Bacillota (17 MAGs from Enterococcaceae, Lactobacillaceae, and Streptococcaceae) and Fusobacteriota (3 MAGs from Fusobacteriaceae).

**Figure 2.**
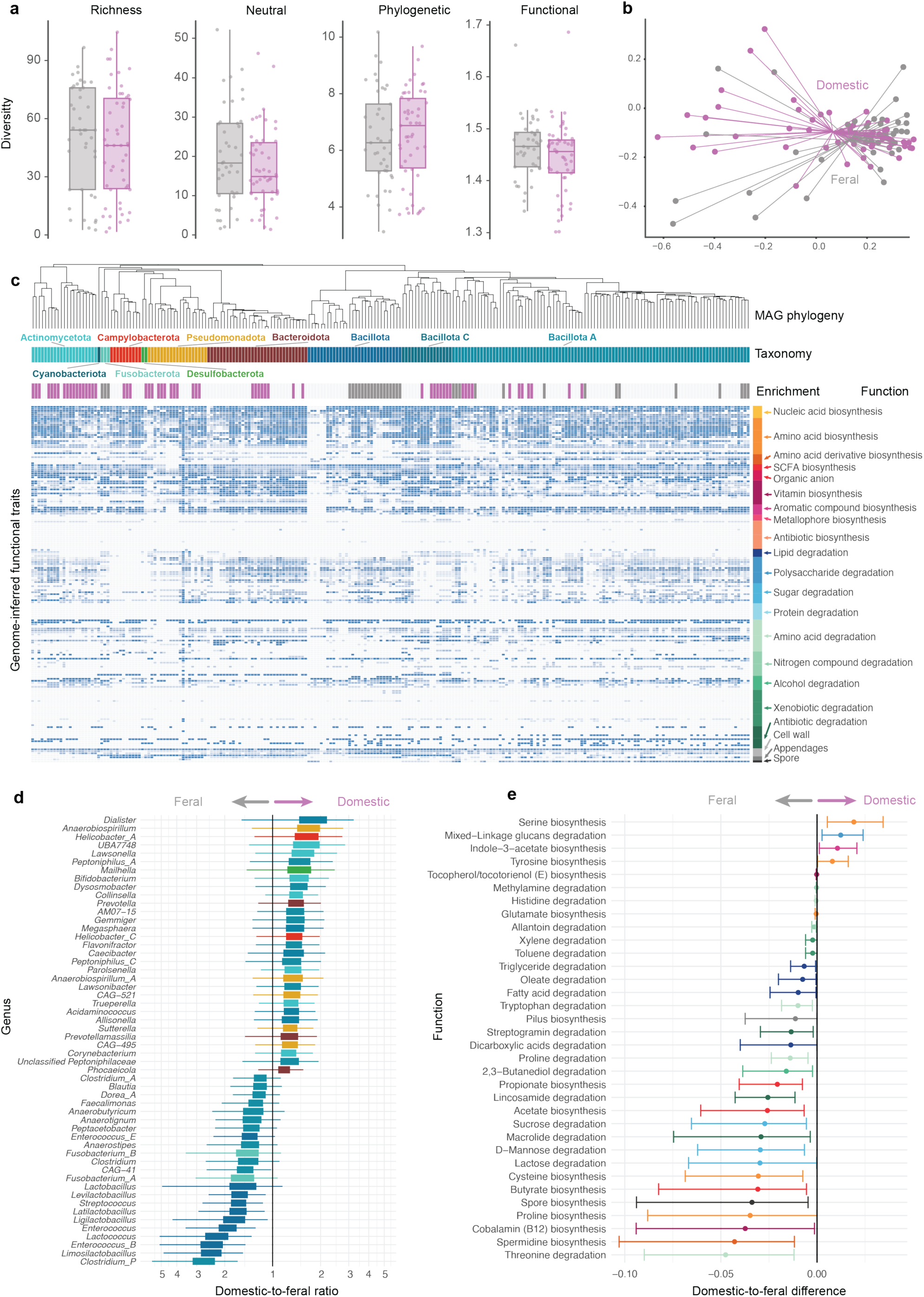
Microbiota differences between domestic and feral cats. **a)** Alpha diversity differences between domestic and feral cats, based on four different Hill numbers diversity metrics. **b)** Compositional differences between domestic and feral cats, samples coloured by domestic or feral origin. **c)** Heatmap of 170 genome-inferred functional traits, with blue colour indicating higher capacity to perform a given metabolic function. Metagenome-assembled genomes (MAGs) are phylogenetically sorted, with their respective enrichment levels annotated above the heatmap. GIFTs are sorted according to their functional group. **d)** Enrichment of bacterial genera towards domestic or feral cats, sorted by effect size and coloured by phyla. **e)** Enrichment of functions towards domestic or feral cats, sorted by effect size and coloured by functional group (see legend in panel c).

Using our HMSC model, we predicted the typical microbial community compositions for feral and domestic cats, while controlling for the rest of the variables. This enabled us to estimate the community-level functional differences of the microbiota associated with both conditions (Figure 2c-e). The microbial community of domestic cats exhibited a higher capacity to degrade polysaccharides such as mixed-linkage glucans. The serine and tyrosine amino acid biosynthetic capacity was also higher in domestic cats, and the capacity to biosynthesise the aromatic compound indole-3-acetate, a major plant growth hormone. The overall repertoire of enriched functions was considerably wider in the case of feral cats (Figure 2e). These included an enhanced capacity for biosynthesising the amino acids glutamate, cysteine and proline, and the amino acid derivative spermidine. The capacities to produce the SCFA acetate, propionate and butyrate, and vitamin E (Tocopherol/tocotorienol) and B12 were also increased. The catabolic repertoire of enriched functions listed capacities for degrading the amino acids histidine, tryptophan, proline, and threonine, the lipids triglyceride, oleate, fatty acid, and dicarboxylic acids, the antibiotics streptogramin, lincosamide and macrolide, the sugar sucrose, D-mannose and lactose, various nitrogen compounds such as methylamine, and allantoin, the alcohol 2,3,-Butanediol and two xenobiotics, such as toluene, and xylene.

## Discussion

Our global genome-resolved metagenomic data set provides unique insights into the taxonomic and functional diversity of cat-associated microbiomes across different geographies and lifestyles. We reconstructed 229 bacterial genomes, with 42 (18%) displaying ANI values below 95% compared to the closest representatives in the GTDB database, indicating potential new species (Rodriguez-R et al., 2024). The gut microbiome of cats was predominantly composed of Bacteroidota, Bacillota A, and Actinomycetota, with notable presences of Campylobacterota and Pseudomonadota, reflecting a typical microbiome profile of a carnivorous mammal (Milani et al., 2020; Zoelzer et al., 2021).

### Cat microbiome variation across geographic space responds to anthropisation rather than climatic variables

Nearly one-third of the bacteria detected in the faecal samples were present in all six sampled locations, indicating a globally distributed core microbiota spanning eight phyla. The remaining bacteria were found in only one or a few locations, resulting in substantial compositional variation across different geographies, consistent with previous observations in other domestic animals, such as horses (Ang et al., 2022) and dogs (Yarlagadda et al., 2022).

The fact that cats from Brazil, Spain, and Denmark formed a cluster, while those from Aruba, Cabo Verde, and Malaysia showed greater geographic and interindividual variation suggests that geographical microbiome variation is influenced more by the level of urbanisation or anthropisation than by climatic variables or geographical proximity. Similar to patterns observed in other species (Dillard et al., 2022), including humans, animals from industrialised areas had more similar and uniform microbial compositions than cats from less urbanised environments. This could be due to the standardisation of dietary resources and the buffering of environmental fluctuations, such as droughts or extreme temperature variations, provided by anthropic environments (Suzuki and Ley, 2020). Nevertheless, these changes were not reflected in alpha diversity metrics, with Aruba and Cabo Verde exhibiting the most and least diverse microbial communities, respectively, across most of the analysed metrics.

### Domestic and feral cats exhibit a taxonomic footprint of lifestyle origin

Despite the significant differences observed across geographic regions, hierarchical modelling of microbial communities revealed taxonomic differences between domestic and feral cats once these regional variations were accounted for. The number of bacterial taxa enriched in domestic and feral cats was similar, corresponding to the fairly balanced alpha diversities in microbiomes from both origins. However, the taxonomic profiles of these enriched taxa differed. Bacteria belonging to Pseudomonadota, Bacteroidota, Campylobacterota, and Desulfovibriota were exclusively enriched in domestic cats. The genera that exhibited the largest effect sizes were *Dialister*, *Anaerobiospirillium* and *Helicobacter_A*. In contrast, the microbiome of feral cats was characterised by an enrichment of many Bacillota and Fusobacteriota species. While members of phylum Bacillota such as *Limosilactobacillus*, *Lactobacillus* and *Ligilactobacillus* are often employed as probiotics due to their capacity to produce lactic acid, many Fusobacteria are pathogenic to humans (Roberts, 2000; Robinson et al., 2020). The enrichment of *Megaspheaera* and *Prevotella* in domestic cats, along with increased *Clostridium* and *Fusobacterium* and decreased *Bifidobacterium* in feral cats, aligns largely with previous studies evaluating the impact of dietary transitions from a kibble diet high in carbohydrates to a more high-protein raw diet in cats and dogs (Bermingham et al., 2017; Butowski et al., 2019; Hooda et al., 2013; Lubbs et al., 2009). Moreover, *Bifidobacterium* and *Dialister*, enriched in domestic cats in our study, were identified as the most featured genera in obese cats distinguishing them from normal cat microbiota (Ma Xiaolei et al., 2022). These shifts in the microbial community are probably the reflection of the dietary differences between domestic and feral cats.

### Feral cats are enriched for microbial functions compared to domestic cats

Beyond taxonomic differences, our bioinformatic and statistical analyses enabled us to capture differences in the functional repertoires of bacteria associated with domestic and feral cats, shedding light on the pathways potentially affecting the domestication and feralisation processes. While taxonomic enrichment was relatively balanced between domestic and feral cats, functional differences were more pronounced. Of the 170 genome-inferred functional traits (GIFTs) analysed, only four showed consistent patterns of enrichment in domestic cats. The reduced number of enriched functional capacities in domestic animals may be associated with their consumption of commercial cat food, which is highly digestible and supplemented with rapidly absorbed nutrients. This likely leads to less substrate available for bacterial activity, depleting the functional repertoire of microbial communities and potentially diminishing their contribution to host biology.

In contrast, we found 30 GIFTs enriched in feral cats. These functional traits encompassed many biosynthetic and degradation pathways, suggesting a more significant role of gut microbiota in feral cats compared to domestic ones. These findings are consistent with earlier observations that domestic animals tend to harbour simplified microbial communities, exhibiting reduced functional potential compared to their wild counterparts (Alessandri et al., 2019). A meta-analysis of the dietary habits of feral cats confirmed that they are obligate carnivores (Plantinga et al., 2011), consuming a diverse range of prey, including rodents, birds, fish, and insects, along with a significant intake of human garbage (Doherty et al., 2015). It has been estimated that 98% of the daily energy intake derives from crude protein and fat (Plantinga et al., 2011). Accordingly, we observed enrichment of many microbial pathways associated with the degradation of such dietary components. Specifically, the microbiome of feral cats showed an enhanced capacity to break down several amino acids and derivatives, as well as lipids including dicarboxylic acids, fatty acids, oleate, and triglycerides. Although only 2% of the daily energy intake has been estimated to derive from nitrogen-free nutrients (Plantinga et al., 2011), we also found enriched capacities for carbohydrate breakdown, indicated by saccharolytic traits for lactose, D-mannose, and sucrose degradation. We hypothesise that this could be linked to consumption of mixed foods found in human waste (Doherty et al., 2015), which may also explain the enrichment of antibiotic degradation pathways, including those for macrolides, lincosamides, and streptogramins.

### Enriched microbial functions align with cat behaviour

The taxonomic and functional differences observed between microbial communities of domestic and feral cats not only reflect dietary changes but also align with the behavioural distinctions seen between these two groups, particularly the fearful and defensive behaviour feral cats exhibited when in contact with humans, in contrast to the docile behaviour observed in domestic cats. This is consistent with studies showing that microbiota can indeed influence animal behaviour (Kamimura et al., 2024; Sudo et al., 2004; Warda et al., 2019; Watanabe et al., 2021; Zhang et al., 2022). As feralisation and domestication are rapid processes involving drastic changes in an animal’s environment and diet, they may trigger microbiota shifts that could alter the microbiota-gut-brain interaction.

Taxonomic shifts observed in feral cats have previously been associated with similar behavioural differences in animals. For instance, *Lactobacillus* and *Streptococcus*, both enriched in feral cats, have been associated with increased fear and anxiety in dogs, chickens, and horses (Bulmer et al., 2019; Mondo et al., 2020; Puetz et al., 2021). Additionally, *Lactobacillus*, *Dorea* and *Blautia* are also more abundant in the gut microbiomes of aggressive dogs (Kirchoff et al., 2019). While these behavioural traits are generally selected against in domestic animals, they may provide an anti-predatory advantage for feral cats. Conversely, *Bifidobacterium*, which was enriched in domestic cats, is commonly used as a probiotic in animals and has been shown to significantly reduce anxiety (Bercik et al., 2011; Desbonnet et al., 2010).

While most of these examples relied on taxonomic profiling derived from 16S rRNA amplicon sequencing, a major advantage of the genome-resolved metagenomics approach we employed lies in its ability to provide direct functional insights from bacterial genomes reconstructed from metagenomic data. For instance, short-chain fatty acids (SCFAs) stood out among the affected biosynthetic pathways, with capacities for synthesising all three major SCFAs—namely butyrate, acetate, and propionate—being enriched in feral cats. SCFAs are produced by bacteria in the lower intestine from indigestible carbohydrates and peptides. Beyond their significance for animal health (Frost et al., 2014; Goswami et al., 2018; Hu et al., 2022; Rowe et al., 2024), SCFAs could also influence observed behaviours by altering brain function via immune, endocrine, and vagal pathways, directly by crossing the blood-brain barrier, or acting as endogenous ligands (Dalile et al., 2019; Kratsman et al., 2016; Mirzaei et al., 2021; Silva et al., 2020). As an example of their relevance in brain function, butyrate has been used to restore cognitive function in neurodegeneration models (Ferrante et al., 2003; Govindarajan et al., 2011; Ryu et al., 2005), and acetate has been shown to offer protection against cognitive impairment in mice (Erny et al., 2021; Zheng et al., 2021). Acetate can also be incorporated into the glutamate–glutamine transcellular cycle and influence hypothalamic neurotransmission, as well as acting as an epigenetic modulator that could have contributed to gene expression changes during domestication (Anastasiadi et al., 2022; Bélteky et al., 2018; Nätt et al., 2012). In fact, acetate synthesis has been recently found to be enriched in the microbiomes of foxes genetically selected for aggressive behaviour (Puetz et al., 2024), which reinforces the possible link between microbiome functions and behaviour in feral cats.

Additionally, the enhanced capacities for synthesising amino acids and derivatives also hold the potential to influence brain function. The capacity to biosynthesise proline, a dietary factor with a significant impact on psychotic disorders (Mayneris-Perxachs et al., 2022; Savio et al., 2012; Yao and Han, 2022), was among the most significantly enriched functions in feral cats. Similarly, the amino acid derivative spermidine has been recently shown to improve cognitive function (Flory et al., 2023; Schroeder et al., 2021). Higher intestinal bacterial degradation of threonine and tryptophan can partially reduce their availability in the brain (Pardridge, 1979; Tovar et al., 1988) thereby influencing the production of key neurotransmitters such as glycine and serotonin, which have important brain functions (Boehm et al., 1998; Gao et al., 2020). Finally, the ability to synthesise cobalamin (vitamin B12) exhibited one of the most significant enrichments in feral over domestic cats. Commonly supplemented in commercial diets, Vitamin B12 can modulate behaviour, its deficiency leading to cognitive impairment and a range of neurological disorders (Kang et al., 2024).

### Insights into the role of microbiomes in feralisation and domestication processes

Feralisation is often recognised as the inverse process of domestication (Price, 1984), where many changes induced by domestication are expected to revert (Henriksen et al., 2018). Despite the broad geographical range screened, our results consistently showed that feral cats gained more microbial functions than their domestic counterparts, confirming our prediction that feral cats would exhibit an enriched functional repertoire compared to domestic cats. Community-level multivariate modelling unveiled significant functional changes in the absence of broad-resolution microbiota markers such as diversity metrics, highlighting the value of statistical tools such as Hmsc for functional microbiome research (Koziol et al., 2023). Although without faecal microbiota transplants and controlled behavioural experiments, we cannot determine causal relationships between microbiota, behaviour and feralisation (Moeller and Sanders, 2020), we conclude that our results point towards a favourable contribution of microbiome changes in the acquisition of physiological and behavioural traits beneficial for a feral lifestyle, as well as a likely effect on the increased aggressiveness and elusiveness observed in the sampled feral cats.

## Materials and Methods

### Sample collection

Veterinary clinics and shelters with trap-neuter-return programmes from six geographically separated countries (Aruba, Brazil, Cabo Verde, Denmark, Malaysia, and Spain) participated in this study. In each location, faecal samples of cats defined as feral (i.e., individuals that had been originally captured in the streets/nature by the Trap-Neuter-Return type programmes) and domesticated (i.e., individuals living with humans prior to entering the clinic or shelter) were collected. To ensure standardised faecal sample collections, sampling kits were sent to each collaborator, including sampling instructions, cat metadata sheets, FTA cards (Whatman FTA Classic), sterile swabs, plastic bags and desiccant packs. The collaborator collected a fresh scat using a sterile swab to transfer some faecal material onto the FTA card, which was then left to dry. Two circles on the filter paper FTA card were collected for each cat faecal sample. The dry FTA card was then stored in a plastic bag until all samples were collected. To characterise the behaviour of the cats categorised as domestic and feral, information about the cats’ behavioural traits towards humans in each facility such as bites, hissing, retreating from, or avoiding human contact was recorded. Cats’ age, sex, date of entry to the facility, date of sampling, and use of antibiotics in the animal were also noted. Once all samples were collected, they were shipped to Denmark for storage at room temperature along with a new desiccant pack.

### DNA extraction

DNA was extracted from 8 feral and 8 domesticated cats from each sampling site following (Bolt Botnen et al., 2023). Briefly, approximately 1/4th of a circle was cut out by a scalpel and placed in a Monarch spin column. 175 µl of elution buffer was added to each spin column before incubating at 37°C for 1 hour. After incubation, each tube was centrifuged at 13,000 x g for 10 minutes. The supernatant was placed in a new Eppendorf tube, ready for storage at −20°C or bead purification with SPRI beads. Bead purification followed the AMPure protocol for a DNA:Bead ratio of 1:1.4 and eluted in 50 μl of elution buffer. DNA was fragmented to an average length of 400 nucleotides using a Covaris M220 ultrasonicator with 50 μl DNA in the microTUBE-50 (Peal power 30, duty factor 20, cycles per burst 50, treatment time 60 seconds).

### Library building

DNA extracts were quantified using a Qubit and then diluted to approximately 100 ng in 16 μl for the library building. Libraries were built using the BEST ligation-based library preparation protocol (Carøe et al., 2018). Per reaction, the end repair master mix contained 2 μl T4 DNA ligase buffer, 1 μl reaction enhancer, 0.5 μl T4 PNK, 0.2 μl T4 polymerase, and 0.2 μl dNTP 25 mM. 3.9 μl of the master mix was added to each sample before incubating at 20℃ for 30 minutes and 65℃ for 30 minutes. The ligation step master mix contained 3 μl PEG 4000 50%, 0.5 μl T4 DNA ligase buffer, and 0.5 μl T4 DNA ligase, per reaction. To each sample 1 μl adapter (50 µM) was added followed by 4 μl of the ligation master mix. The samples were then incubated at 20°C for 30 minutes and 65°C for 10 minutes. Fill-in master mix contained, per reaction, 3 μl ddH_2_O, 1 μl Isothermal buffer, 0.2 μl dNTP 25 mM, and 0.8 μl Bst 2.0 polymerase. 5 μl of fill-in master mix was added to each sample followed by incubation at 65℃ for 15 min and 80℃ for 15 minutes. Each library was then diluted 1:2 with molecular-grade water. Bead purification followed the AMPure purification protocol, substituted with SPRI beads, at a ratio of 1:1.4 and eluted in 30 μl of elution buffer.

Dilutions (1:100) of each library were prepared for qPCR. A qPCR master mix was prepared with 11.8 μl ddH_2_O, 2.5 μl 10x buffer, 2.5 μl MgCl_2_ 25mM, 0.5 μl BSA, 0.5 μl SYBR, 0.2 μl dNTP 25 mM, 0.5 forward primer (BGI-F 10 μm) and 0.5 μl TaqGold, per reaction. Each reaction consisted of 10 μl diluted DNA, 1 μl reverse primer (BGI-R 10 μM) and 19 μl master mix. qPCR settings were as follows: 95°C for 10 minutes, 30 cycles of 95°C for 20 seconds, 60°C for 30 seconds, 72°C for 40 seconds, followed by a final 72°C for 7 minutes and a 4°C hold. The cycle threshold determined the number of cycles required for indexing. For indexing, a master mix was prepared with 12.3 μl ddH_2_O, 2.5 μl 10x buffer, 2.5 μl MgCl_2_ 25mM, 0.5 μl BSA, 0.2 μl dNTP 25 mM, and 0.5 μl TaqGold, per reaction. Each reaction consisted of 10 μl undiluted DNA, 1 μl of each indexed primer (BGI-R and BGI-F 10 μM) and 19 μl master mix. PCR settings were as follows: 95°C for 10 minutes, 7-11 cycles of 95°C for 20 seconds, 60°C for 30 seconds, 72°C for 40 seconds, followed by a final 72°C for 7 minutes and a 4°C hold. Libraries were then bead purified with a ratio of 1:1.4 and eluted in 30 μl elution buffer and then quantified on the Qubit and 2-3 μl aliquoted for fragment analysis. Libraries were pooled together based on country. Libraries from five locations were sequenced on a DNBseq platform at BGI Denmark, while one location (Denmark) was sequenced using Illumina technology at a NovaSeq 6000 platform at Novogene (UK), after ensuring that both platforms are compatible for combined microbiome analyses (Mak et al., 2017; Smith et al., 2019).

### Bioinformatics

Using a custom snakemake-based (Mölder et al., 2021) pipeline, paired-end reads were trimmed and quality controlled using fastp v0.20.1 (Chen et al., 2018), with the following options: —trim_poly_g, — trim_poly_x, —n_base_limit 5, —qualified_quality_phred 20, —length_required 35. Processed reads were then mapped to the concatenated host reference genome assemblies (Human; GRCh38.p13), Cat; F.catus_Fca126_mat1.0) using Bowtie2 (Langmead and Salzberg, 2012) and samtools (Li et al., 2009) (default settings). Unaligned reads from each geographic location were then pooled and coassembled using metaSPAdes (Nurk et al., 2017) with the following kmer sizes: 21,29,39,59,79,99,119. Coassembled contigs shorter than 1,500 bp were removed. Each sample’s reads were then mapped its appropriate coassembly using Bowtie2, and the resulting BAMs were used as input to MetaWRAP’s (Uritskiy et al., 2018) binning module, and the coassembled contigs binned using CONCOCT (Alneberg et al., 2014), MaxBin2 (Wu et al., 2016), and MetaBAT2 (Kang et al., 2019). The output bins were automatically refined using MetaWRAP’s bin_refinement module, with a minimum CheckM (Parks et al., 2015) completeness score of 70%, and a minimum checkM contamination score of 10%. dRep (Olm et al., 2017) was then used to dereplicate the refined bins, first into primary clusters >90% average nucleotide identity (ANI) using MASH (Ondov et al., 2016), then into secondary clusters >98% ANI with ANImf (Kurtz et al., 2004; Richter and Rosselló-Móra, 2009). The non-host quality-controlled reads were then mapped against this dereplicated MAG catalogue as above. The resulting BAMs were then profiled using CoverM (https://github.com/wwood/CoverM) to create the final sample count table. The dereplicated MAGs were taxonomically annotated using the GTDB-tk (Chaumeil et al., 2019; Parks et al., 2020), which uses pplacer (Matsen et al., 2010), prodigal (Hyatt et al., 2010), HMMER3 (Eddy, 2011), FastANI (Jain et al., 2018), and FastTree (Price et al., 2010). The MAGs were also functionally annotated using the DRAM pipeline (Shaffer et al., 2020), which searches predicted proteins against multiple databases (El-Gebali et al., 2019; Kanehisa et al., 2017; Rawlings et al., 2010; Suzek et al., 2015) using MMseqs2 (Steinegger and Söding, 2017). Finally, gene annotations were distilled into Genome-Inferred Functional Traits (GIFTs) using distillR (https://github.com/anttonalberdi/distillR), producing biologically meaningful annotations that highlight each bacterial genome’s potential to degrade or synthesise compounds relevant to host metabolism. DistillR utilises a curated database of over 300 metabolic pathways, employing KEGG and Enzyme Commission (EC) identifiers to calculate standardised GIFT values. These values range from 0 to 1, where 0 signifies the absence of all genes associated with a particular metabolic pathway, and 1 indicates the presence of all necessary genes. For example, if a pathway step requires two specific identifiers, it is deemed complete when both are present, half-complete when only one is present, and empty if neither is present.

### Statistics

#### Behavioural phenotypes in domestic and feral cats

We follow the earlier definitions of feral cats (Crowley et al., 2020; Gosling et al., 2013), defining a feral cat as an unowned specimen capable of surviving with or without direct human intervention, and additionally showing fearful or defensive behaviour upon human contact. In each location, cats captured in the streets or nature were classified as feral, while those living in a residence with their owners were classified as domestic. To ensure the origin of the cats correlated with the behavioural phenotype, we analysed the differences in cats’ fearfulness or defensive behaviour towards humans using chi-tests.

#### Alpha and beta diversities

Alpha and beta diversity analyses were performed within the Hill numbers framework (Alberdi and Gilbert, 2019), using different combinations of orders of diversity (q) and regularity in Hilldiv2 R package (https://github.com/anttonalberdi/Hilldiv2). ‘Richness’ refers to the neutral Hill number of q=0, which is limited to the counts of MAGs. ‘Neutral’ refers to the neutral Hill number of q=1, in which the MAGs are weighted according to their relative representation. In the case of alpha diversity, it equals the exponential of the Shannon index, also known as Shannon diversity. ‘Phylogenetic’ refers to the phylogenetic Hill number of q=1, which accounts for the phylogenetic relationships between MAGs derived from the phylogenetic tree. ‘Functional’ refers to the functional Hill number of q=1, which accounts for the functional distances between MAGs derived from the GIFT table. Compositional differences were quantified using pairwise dissimilarity estimation via the Jaccard-type turnover metric (S) derived from the Hill numbers beta diversity. All four metrics were employed for alpha and beta diversity comparisons. Differences in alpha diversity between locations were assessed using linear models, while differences between origins were tested using linear mixed models with location as a random effect. PERMANOVA analysis was performed to compare the beta diversity differences between locations and origins using vegan::adonis2 function.

#### Hierarchical modelling of species communities

To understand taxonomic and functional responses to the analysed variables, we built models of species communities using Hmsc, a multivariate hierarchical generalised linear model that uses Bayesian inference (Tikhonov et al., 2020). In the species matrix Y, which typically includes species abundance or occurrence values, we included the genome-size normalised and log-transformed counts of MAGs. In the environmental variable matrix X, we included the categorical variables origin and sex, as well as the log-transformed sequencing depth varying sequencing efforts across samples. Geographic location was included as a random effect to account for the non-independence of samples collected from the same location. Additionally, we included the MAG phylogeny in the matrix C to quantify the degree of phylogenetic signal in the MAG’s responses to the variables.

We fitted the models assuming default priors and sampled the posterior distribution running four Markov Chain Monte Carlo (MCMC) chains, each running for 3,750 iterations with 1,250 discarded as burn-in. We thinned by 10 to obtain a total of 250 posterior samples per chain and 1,000 total posterior samples. To test for MCMC convergence we measured the potential scale reduction factor for the beta (response to variables) and rho parameters (phylogenetic signal). We then performed five-fold cross-validation to evaluate the predictive performance of the models for the analysed MAGs in terms of R^2^. Once the models were fitted, we quantified the posterior estimates for the variable to origin to identify MAGs were considered significantly associated with either feral or domestic origins when 90% of the posterior credible interval of the beta parameter measuring the effect of origin was either positive or negative (i.e. using a significance threshold of 0.9 posterior probability). To estimate functional differences between microbial communities associated with feral or domestic origins, we compared the community-level functional attributes of the subset of MAGs with significant association with origin. To minimise the risk of false positives, we filtered out MAGs with low predictive capacity related to origin. This filtering was based on a metric calculated as the product of the variance attributed to origin (derived from variance partitioning) and the R² value obtained from five-fold cross-validation.

All statistical analyses performed in this study are fully reproduced in a Rmarkdown webbook (https://alberdilab.github.io/domestic_feral_cat_metagenomics).

## Data accessibility

Raw metagenomic sequences have been deposited under the BioProject accession PRJEB79769.

## Acknowledgements

We would like to thank Pieter Barendsen, DVM, Aruba Vets Wayaca (Aruba), Flávia P. Tirelli, Ph.D, Pontifícia Universidade Católica do Rio Grande do Sul (Porto Alegre, Brazil), Prof. Augusto Faustino, University of Porto (Portugal, Cabo Verde), Therese Wilbert, DVM (Denmark), Kattens Værn, Brøndby (Denmark), Gabriel Bustillo, Fundación Protectora de Animales del Principado de Asturias (Spain), and Dr. Natasha Lee, DVM (Malaysia) for their efforts in sample collection. This work was supported by the Villum Experiment under grant 17417, the Carlsberg Foundation under Grant CF20-0460 and the Danish National Research Foundation under Grant DNRF143 “A Center for Evolutionary Hologenomics”.

## References

Alberdi A, Aizpurua O, Bohmann K, Zepeda-Mendoza ML, Gilbert MTP. 2016. Do Vertebrate Gut Metagenomes Confer Rapid Ecological Adaptation? Trends Ecol Evol 31:689–699.

Alberdi A, Gilbert MTP. 2019. A guide to the application of Hill numbers to DNA-based diversity analyses. Mol Ecol Resour 19:804–817.

Alessandri G, Milani C, Mancabelli L, Mangifesta M, Lugli GA, Viappiani A, Duranti S, Turroni F, Ossiprandi MC, van Sinderen D, Ventura M. 2019. The impact of human-facilitated selection on the gut microbiota of domesticated mammals. FEMS Microbiol Ecol 95. doi:10.1093/femsec/fiz121

Alneberg J, Bjarnason BS, de Bruijn I, Schirmer M, Quick J, Ijaz UZ, Lahti L, Loman NJ, Andersson AF, Quince C. 2014. Binning metagenomic contigs by coverage and composition. Nat Methods 11:1144–1146.

Anastasiadi D, Piferrer F, Wellenreuther M, Benítez Burraco A. 2022. Fish as model systems to study epigenetic drivers in human self-domestication and neurodevelopmental cognitive disorders. Genes (Basel) 13:987.

Ang L, Vinderola G, Endo A, Kantanen J, Jingfeng C, Binetti A, Burns P, Qingmiao S, Suying D, Zujiang Y, Rios-Covian D, Mantziari A, Beasley S, Gomez-Gallego C, Gueimonde M, Salminen S. 2022. Gut Microbiome Characteristics in feral and domesticated horses from different geographic locations. Commun Biol 5:172.

Baca M, Popović D, Panagiotopoulou H, Marciszak A, Krajcarz M, Krajcarz MT, Makowiecki D, Węgleński P, Nadachowski A. 2018. Human-mediated dispersal of cats in the Neolithic Central Europe. Heredity 121:557–563.

Bélteky J, Agnvall B, Bektic L, Höglund A, Jensen P, Guerrero-Bosagna C. 2018. Epigenetics and early domestication: differences in hypothalamic DNA methylation between red junglefowl divergently selected for high or low fear of humans. Genet Sel Evol 50:13.

Bercik P, Park AJ, Sinclair D, Khoshdel A, Lu J, Huang X, Deng Y, Blennerhassett PA, Fahnestock M, Moine D, Berger B, Huizinga JD, Kunze W, McLean PG, Bergonzelli GE, Collins SM, Verdu EF. 2011. The anxiolytic effect of Bifidobacterium longum NCC3001 involves vagal pathways for gut-brain communication. Neurogastroenterol Motil 23:1132–1139.

Bermingham EN, Maclean P, Thomas DG, Cave NJ, Young W. 2017. Key bacterial families (Clostridiaceae, Erysipelotrichaceae and Bacteroidaceae) are related to the digestion of protein and energy in dogs. PeerJ 5:e3019.

Boehm G, Cervantes H, Georgi G, Jelinek J, Sawatzki G, Wermuth B, Colombo JP. 1998. Effect of increasing dietary threonine intakes on amino acid metabolism of the central nervous system and peripheral tissues in growing rats. Pediatr Res 44:900–906.

Bolt Botnen A, Bjørnsen MB, Alberdi A, Gilbert MTP, Aizpurua O. 2023. A simplified protocol for DNA extraction from FTA cards for faecal microbiome studies. Heliyon 9:e12861.

Bonacic C, Almuna R, Ibarra JT. 2019. Biodiversity Conservation Requires Management of Feral Domestic Animals. Trends Ecol Evol 34:683–686.

Bradshaw JWS. 2006. The evolutionary basis for the feeding behavior of domestic dogs (Canis familiaris) and cats (Felis catus). J Nutr 136:1927S–1931S.

Bradshaw JWS, Horsfield GF, Allen JA, Robinson IH. 1999. Feral cats: their role in the population dynamics of Felis catus. Appl Anim Behav Sci 65:273–283.

Bulmer LS, Murray J-A, Burns NM, Garber A, Wemelsfelder F, McEwan NR, Hastie PM. 2019. High-starch diets alter equine faecal microbiota and increase behavioural reactivity. Sci Rep 9:18621.

Butowski CF, Thomas DG, Young W, Cave NJ, McKenzie CM, Rosendale DI, Bermingham EN. 2019. Addition of plant dietary fibre to a raw red meat high protein, high fat diet, alters the faecal bacteriome and organic acid profiles of the domestic cat (Felis catus). PLoS One 14:e0216072.

Candela M, Biagi E, Maccaferri S, Turroni S, Brigidi P. 2012. Intestinal microbiota is a plastic factor responding to environmental changes. Trends Microbiol 20:385–391.

Carøe C, Gopalakrishnan S, Vinner L, Mak SST, Sinding MHS, Samaniego JA, Wales N, Sicheritz-Pontén T, Gilbert MTP. 2018. Single-tube library preparation for degraded DNA. Methods Ecol Evol 9:410–419.

Cecchetti M, Crowley SL, McDonald RA. 2021. Drivers and facilitators of hunting behaviour in domestic cats and options for management. Mamm Rev 51:307–322.

Chaumeil P-A, Mussig AJ, Hugenholtz P, Parks DH. 2019. GTDB-Tk: a toolkit to classify genomes with the Genome Taxonomy Database. Bioinformatics. doi:10.1093/bioinformatics/btz848

Chen S, Zhou Y, Chen Y, Gu J. 2018. fastp: an ultra-fast all-in-one FASTQ preprocessor. Bioinformatics 34:i884–i890.

Clutton-Brock J. 1990. Α Natural History of Domesticated Mammals. Praehistorische Zeitschrift 65:73–76.

Crowley SL, Cecchetti M, McDonald RA. 2020. Our Wild Companions: Domestic cats in the Anthropocene. Trends Ecol Evol 35:477–483.

Dalile B, Van Oudenhove L, Vervliet B, Verbeke K. 2019. The role of short-chain fatty acids in microbiota-gut-brain communication. Nat Rev Gastroenterol Hepatol 16:461–478.

Desbonnet L, Garrett L, Clarke G, Kiely B, Cryan JF, Dinan TG. 2010. Effects of the probiotic Bifidobacterium infantis in the maternal separation model of depression. Neuroscience 170:1179–1188.

Dillard BA, Chung AK, Gunderson AR, Campbell-Staton SC, Moeller AH. 2022. Humanization of wildlife gut microbiota in urban environments. Elife 11. doi:10.7554/eLife.76381

Doherty TS, Davis RA, van Etten EJB, Algar D, Collier N, Dickman CR, Edwards G, Masters P, Palmer R, Robinson S. 2015. A continental-scale analysis of feral cat diet in Australia. J Biogeogr 42:964–975.

Driscoll CA, Menotti-Raymond M, Roca AL, Hupe K, Johnson WE, Geffen E, Harley EH, Delibes M, Pontier D, Kitchener AC, Yamaguchi N, O’brien SJ, Macdonald DW. 2007. The Near Eastern origin of cat domestication. Science 317:519–523.

Dueñas M-A, Hemming DJ, Roberts A, Diaz-Soltero H. 2021. The threat of invasive species to IUCN-listed critically endangered species: A systematic review. Global Ecology and Conservation 26:e01476.

Eddy SR. 2011. Accelerated Profile HMM Searches. PLoS Comput Biol 7:e1002195.

El-Gebali S, Mistry J, Bateman A, Eddy SR, Luciani A, Potter SC, Qureshi M, Richardson LJ, Salazar GA, Smart A, Sonnhammer ELL, Hirsh L, Paladin L, Piovesan D, Tosatto SCE, Finn RD. 2019. The Pfam protein families database in 2019. Nucleic Acids Res 47:D427–D432.

Erny D, Dokalis N, Mezö C, Castoldi A, Mossad O, Staszewski O, Frosch M, Villa M, Fuchs V, Mayer A, Neuber J, Sosat J, Tholen S, Schilling O, Vlachos A, Blank T, Gomez de Agüero M, Macpherson AJ, Pearce EJ, Prinz M. 2021. Microbiota-derived acetate enables the metabolic fitness of the brain innate immune system during health and disease. Cell Metab 33:2260–2276.e7.

Ferrante RJ, Kubilus JK, Lee J, Ryu H, Beesen A, Zucker B, Smith K, Kowall NW, Ratan RR, Luthi-Carter R, Hersch SM. 2003. Histone deacetylase inhibition by sodium butyrate chemotherapy ameliorates the neurodegenerative phenotype in Huntington’s disease mice. J Neurosci 23:9418–9427.

Flory M, Gao A, Morrow M, Alam A. 2023. Colonic spermidine promotes proliferation and migration of intestinal epithelial cells. bioRxiv. doi:10.1101/2023.09.26.559404

Forsythe P, Bienenstock J, Kunze WA. 2014. Vagal pathways for microbiome-brain-gut axis communication. Adv Exp Med Biol 817:115–133.

Frost G, Sleeth ML, Sahuri-Arisoylu M, Lizarbe B, Cerdan S, Brody L, Anastasovska J, Ghourab S, Hankir M, Zhang S, Carling D, Swann JR, Gibson G, Viardot A, Morrison D, Louise Thomas E, Bell JD. 2014. The short-chain fatty acid acetate reduces appetite via a central homeostatic mechanism. Nat Commun 5:3611.

Gao K, Mu C-L, Farzi A, Zhu W-Y. 2020. Tryptophan Metabolism: A Link Between the Gut Microbiota and Brain. Adv Nutr 11:709–723.

Gering E, Incorvaia D, Henriksen R, Conner J, Getty T, Wright D. 2019. Getting Back to Nature: Feralization in Animals and Plants. Trends Ecol Evol 34:1137–1151.

Gosling L, Stavisky J, Dean R. 2013. What is a feral cat?: Variation in definitions may be associated with different management strategies. J Feline Med Surg 15:759–764.

Goswami C, Iwasaki Y, Yada T. 2018. Short-chain fatty acids suppress food intake by activating vagal afferent neurons. J Nutr Biochem 57:130–135.

Govindarajan N, Agis-Balboa RC, Walter J, Sananbenesi F, Fischer A. 2011. Sodium butyrate improves memory function in an Alzheimer’s disease mouse model when administered at an advanced stage of disease progression. Journal of Alzheimer’s Disease 26:187–197.

Gross M. 2020. Of mice and men, cats and grains. Curr Biol 30:R783–R786.

Henriksen R, Gering E, Wright D. 2018. Feralisation—The Understudied Counterpoint to Domestication In: Pontarotti P, editor. Origin and Evolution of Biodiversity. Cham: Springer International Publishing. pp. 183–195.

Henry LP, Bruijning M, Forsberg SKG, Ayroles JF. 2021. The microbiome extends host evolutionary potential. Nat Commun 12:5141.

Hooda S, Vester Boler BM, Kerr KR, Dowd SE, Swanson KS. 2013. The gut microbiome of kittens is affected by dietary protein:carbohydrate ratio and associated with blood metabolite and hormone concentrations. Br J Nutr 109:1637–1646.

Hu T, Wu Q, Yao Q, Jiang K, Yu J, Tang Q. 2022. Short-chain fatty acid metabolism and multiple effects on cardiovascular diseases. Ageing Res Rev 81:101706.

Hyatt D, Chen G-L, Locascio PF, Land ML, Larimer FW, Hauser LJ. 2010. Prodigal: prokaryotic gene recognition and translation initiation site identification. BMC Bioinformatics 11:119.

Jain C, Rodriguez-R LM, Phillippy AM, Konstantinidis KT, Aluru S. 2018. High throughput ANI analysis of 90K prokaryotic genomes reveals clear species boundaries. Nat Commun 9:5114.

Johnson KV-A, Foster KR. 2018. Why does the microbiome affect behaviour? Nat Rev Microbiol 16:647–655.

Kamimura I, Miyauchi E, Takeuchi T, Tsuchiya N, Tamura K, Uesugi A, Negishi H, Taida T, Kato T, Kawasumi M, Nagasawa M, Mogi K, Ohno H, Kikusui T. 2024. Modulation of gut microbiota composition due to early weaning stress induces depressive behavior during the juvenile period in mice. Anim Microbiome 6:33.

Kanehisa M, Furumichi M, Tanabe M, Sato Y, Morishima K. 2017. KEGG: new perspectives on genomes, pathways, diseases and drugs. Nucleic Acids Res 45:D353–D361.

Kang DD, Li F, Kirton E, Thomas A, Egan R, An H, Wang Z. 2019. MetaBAT 2: an adaptive binning algorithm for robust and efficient genome reconstruction from metagenome assemblies. PeerJ 7:e7359.

Kang WK, Florman JT, Araya A, Fox BW, Thackeray A, Schroeder FC, Walhout AJM, Alkema MJ. 2024. Vitamin B12 produced by gut bacteria modulates cholinergic signalling. Nat Cell Biol 26:72–85.

Kirchoff NS, Udell MAR, Sharpton TJ. 2019. The gut microbiome correlates with conspecific aggression in a small population of rescued dogs (Canis familiaris). PeerJ 7:e6103.

Koziol A, Odriozola I, Leonard A, Eisenhofer R, San José C, Aizpurua O, Alberdi A. 2023. Mammals show distinct functional gut microbiome dynamics to identical series of environmental stressors. MBio 14:e0160623.

Kratsman N, Getselter D, Elliott E. 2016. Sodium butyrate attenuates social behavior deficits and modifies the transcription of inhibitory/excitatory genes in the frontal cortex of an autism model. Neuropharmacology 102:136–145.

Kurtz S, Phillippy A, Delcher AL, Smoot M, Shumway M, Antonescu C, Salzberg SL. 2004. Versatile and open software for comparing large genomes. Genome Biol 5:R12.

Langmead B, Salzberg SL. 2012. Fast gapped-read alignment with Bowtie 2. Nat Methods 9:357–359.

Lee W-J, Hase K. 2014. Gut microbiota-generated metabolites in animal health and disease. Nat Chem Biol 10:416–424.

Levy JK, Gale DW, Gale LA. 2003. Evaluation of the effect of a long-term trap-neuter-return and adoption program on a free-roaming cat population. J Am Vet Med Assoc 222:42–46.

Li H, Handsaker B, Wysoker A, Fennell T, Ruan J, Homer N, Marth G, Abecasis G, Durbin R, 1000 Genome Project Data Processing Subgroup. 2009. The Sequence Alignment/Map format and SAMtools. Bioinformatics 25:2078–2079.

Lubbs DC, Vester BM, Fastinger ND, Swanson KS. 2009. Dietary protein concentration affects intestinal microbiota of adult cats: a study using DGGE and qPCR to evaluate differences in microbial populations in the feline gastrointestinal tract. J Anim Physiol Anim Nutr 93:113–121.

Mak SST, Gopalakrishnan S, Carøe C, Geng C, Liu S, Sinding M-HS, Kuderna LFK, Zhang W, Fu S, Vieira FG, Germonpré M, Bocherens H, Fedorov S, Petersen B, Sicheritz-Pontén T, Marques-Bonet T, Zhang G, Jiang H, Gilbert MTP. 2017. Comparative performance of the BGISEQ-500 vs Illumina HiSeq2500 sequencing platforms for palaeogenomic sequencing. Gigascience 6:1–13.

Ma L-C, Zhao H-Q, Wu LB, Cheng Z-L, Liu C. 2023. Impact of the microbiome on human, animal, and environmental health from a One Health perspective. Science in One Health 2:100037.

Matsen FA, Kodner RB, Armbrust EV. 2010. pplacer: linear time maximum-likelihood and Bayesian phylogenetic placement of sequences onto a fixed reference tree. BMC Bioinformatics 11:538.

Ma Xiaolei, Brinker Emily, Graff Emily C., Cao Wenqi, Gross Amanda L., Johnson Aime K., Zhang Chao, Martin Douglas R., Wang Xu. 2022. Whole-Genome Shotgun Metagenomic Sequencing Reveals Distinct Gut Microbiome Signatures of Obese Cats. Microbiology Spectrum 10:e00837–22.

Mayer EA. 2011. Gut feelings: the emerging biology of gut–brain communication. Nat Rev Neurosci 12:453–466.

Mayneris-Perxachs J, Castells-Nobau A, Arnoriaga-Rodríguez M, Martin M, de la Vega-Correa L, Zapata C, Burokas A, Blasco G, Coll C, Escrichs A, Biarnés C, Moreno-Navarrete JM, Puig J, Garre-Olmo J, Ramos R, Pedraza S, Brugada R, Vilanova JC, Serena J, Gich J, Ramió-Torrentà L, Pérez-Brocal V, Moya A, Pamplona R, Sol J, Jové M, Ricart W, Portero-Otin M, Deco G, Maldonado R, Fernández-Real JM. 2022. Microbiota alterations in proline metabolism impact depression. Cell Metab 34:681–701.e10.

Medina FM, Bonnaud E, Vidal E, Tershy BR, Zavaleta ES, Josh Donlan C, Keitt BS, Corre M, Horwath SV, Nogales M. 2011. A global review of the impacts of invasive cats on island endangered vertebrates. Glob Chang Biol 17:3503–3510.

Milani C, Alessandri G, Mancabelli L, Mangifesta M, Lugli GA, Viappiani A, Longhi G, Anzalone R, Duranti S, Turroni F, Ossiprandi MC, van Sinderen D, Ventura M. 2020. Multi-omics Approaches To Decipher the Impact of Diet and Host Physiology on the Mammalian Gut Microbiome. Appl Environ Microbiol 86. doi:10.1128/AEM.01864-20

Mirzaei R, Bouzari B, Hosseini-Fard SR, Mazaheri M, Ahmadyousefi Y, Abdi M, Jalalifar S, Karimitabar Z, Teimoori A, Keyvani H, Zamani F, Yousefimashouf R, Karampoor S. 2021. Role of microbiota-derived short-chain fatty acids in nervous system disorders. Biomed Pharmacother 139:111661.

Moeller AH, Sanders JG. 2020. Roles of the gut microbiota in the adaptive evolution of mammalian species. Philos Trans R Soc Lond B Biol Sci 375:20190597.

Mölder F, Jablonski KP, Letcher B, Hall MB, Tomkins-Tinch CH, Sochat V, Forster J, Lee S, Twardziok SO, Kanitz A, Wilm A, Holtgrewe M, Rahmann S, Nahnsen S, Köster J. 2021. Sustainable data analysis with Snakemake. F1000Res 10:33.

Mondo E, Barone M, Soverini M, D’Amico F, Cocchi M, Petrulli C, Mattioli M, Marliani G, Candela M, Accorsi PA. 2020. Gut microbiome structure and adrenocortical activity in dogs with aggressive and phobic behavioral disorders. Heliyon 6:e03311.

Nätt D, Rubin C-J, Wright D, Johnsson M, Beltéky J, Andersson L, Jensen P. 2012. Heritable genome-wide variation of gene expression and promoter methylation between wild and domesticated chickens. BMC Genomics 13:59.

Nurk S, Meleshko D, Korobeynikov A, Pevzner PA. 2017. metaSPAdes: a new versatile metagenomic assembler. Genome Res 27:824–834.

Olm MR, Brown CT, Brooks B, Banfield JF. 2017. dRep: a tool for fast and accurate genomic comparisons that enables improved genome recovery from metagenomes through de-replication. ISME J 11:2864–2868.

Ondov BD, Treangen TJ, Melsted P, Mallonee AB, Bergman NH, Koren S, Phillippy AM. 2016. Mash: fast genome and metagenome distance estimation using MinHash. Genome Biol 17:132.

Ottoni C, Van Neer W. 2020. The Dispersal of the Domestic Cat: Paleogenetic and Zooarcheological Evidence. Near Eastern Archaeology 83:38–45.

Palmas P, Jourdan H, Rigault F, Debar L, De Meringo H, Bourguet E, Mathivet M, Lee M, Adjouhgniope R, Papillon Y, Bonnaud E, Vidal E. 2017. Feral cats threaten the outstanding endemic fauna of the New Caledonia biodiversity hotspot. Biol Conserv 214:250–259.

Pardridge WM. 1979. Tryptophan transport through the blood-brain barrier: in vivo measurement of free and albumin-bound amino acid. Life Sci 25:1519–1528.

Parks DH, Chuvochina M, Chaumeil P-A, Rinke C, Mussig AJ, Hugenholtz P. 2020. A complete domain-to-species taxonomy for Bacteria and Archaea. Nat Biotechnol 38:1079–1086.

Parks DH, Imelfort M, Skennerton CT, Hugenholtz P, Tyson GW. 2015. CheckM: assessing the quality of microbial genomes recovered from isolates, single cells, and metagenomes. Genome Res 25:1043–1055.

Plantinga EA, Bosch G, Hendriks WH. 2011. Estimation of the dietary nutrient profile of free-roaming feral cats: possible implications for nutrition of domestic cats. Br J Nutr 106 Suppl 1:S35–48.

Price EO. 1984. Behavioral Aspects of Animal Domestication. Q Rev Biol 59:1–32.

Price MN, Dehal PS, Arkin AP. 2010. FastTree 2--approximately maximum-likelihood trees for large alignments. PLoS One 5:e9490.

Puetz LC, Delmont TO, Aizpurua O, Guo C, Zhang G, Katajamaa R, Jensen P, Gilbert MTP. 2021. Gut Microbiota linked with reduced fear of humans in red junglefowl has implications for early domestication. Advanced Genetics 2:2100018.

Puetz LC, Delmont TO, Mitchell AL, Finn R, Zhang G, Shepeleva DV, Kharlamova AV, Kukekova A, Trut LN, Gilbert MTP. 2024. Gut microbiome community structure correlates with different behavioral phenotypes in the Belyaev farm-fox experiment. Research Square. doi:10.21203/rs.3.rs-4697888/v1

Rawlings ND, Barrett AJ, Bateman A. 2010. MEROPS: the peptidase database. Nucleic Acids Res 38:D227–33.

Richter M, Rosselló-Móra R. 2009. Shifting the genomic gold standard for the prokaryotic species definition. Proc Natl Acad Sci U S A 106:19126–19131.

Roberts GL. 2000. Fusobacterial infections: an underestimated threat. Br J Biomed Sci 57:156–162.

Robinson A, Wilde J, Allen-Vercoe E. 2020. Fusobacteria: Physiology, form, and functionColorectal Neoplasia and the Colorectal Microbiome. Elsevier. pp. 95–134.

Rodriguez-R LM, Conrad RE, Viver T, Feistel DJ, Lindner BG, Venter SN, Orellana LH, Amann R, Rossello-Mora R, Konstantinidis KT. 2024. An ANI gap within bacterial species that advances the definitions of intra-species units. MBio 15:e0269623.

Rosenberg E, Zilber-Rosenberg I. 2022. Special issue: The role of microorganisms in the evolution of animals and plants. Microorganisms 10:250.

Rowe JC, Winston JA, Parker VJ, McCool KE, Suchodolski JS, Lopes R, Steiner JM, Gilor C, Rudinsky AJ. 2024. Gut microbiota promoting propionic acid production accompanies caloric restriction-induced intentional weight loss in cats. Sci Rep 14:11901.

Ryu H, Smith K, Camelo SI, Carreras I, Lee J, Iglesias AH, Dangond F, Cormier KA, Cudkowicz ME, Brown RH Jr, Ferrante RJ. 2005. Sodium phenylbutyrate prolongs survival and regulates expression of anti-apoptotic genes in transgenic amyotrophic lateral sclerosis mice: Histone deacetylase therapy in amyotrophic lateral sclerosis. J Neurochem 93:1087–1098.

Savio LEB, Vuaden FC, Piato AL, Bonan CD, Wyse ATS. 2012. Behavioral changes induced by long-term proline exposure are reversed by antipsychotics in zebrafish. Prog Neuropsychopharmacol Biol Psychiatry 36:258–263.

Schroeder S, Hofer SJ, Zimmermann A, Pechlaner R, Dammbrueck C, Pendl T, Marcello GM, Pogatschnigg V, Bergmann M, Müller M, Gschiel V, Ristic S, Tadic J, Iwata K, Richter G, Farzi A, Üçal M, Schäfer U, Poglitsch M, Royer P, Mekis R, Agreiter M, Tölle RC, Sótonyi P, Willeit J, Mairhofer B, Niederkofler H, Pallhuber I, Rungger G, Tilg H, Defrancesco M, Marksteiner J, Sinner F, Magnes C, Pieber TR, Holzer P, Kroemer G, Carmona-Gutierrez D, Scorrano L, Dengjel J, Madl T, Sedej S, Sigrist SJ, Rácz B, Kiechl S, Eisenberg T, Madeo F. 2021. Dietary spermidine improves cognitive function. Cell Rep 35:108985.

Shaffer M, Borton MA, McGivern BB, Zayed AA, La Rosa SL, Solden LM, Liu P, Narrowe AB, Rodríguez-Ramos J, Bolduc B, Gazitúa MC, Daly RA, Smith GJ, Vik DR, Pope PB, Sullivan MB, Roux S, Wrighton KC. 2020. DRAM for distilling microbial metabolism to automate the curation of microbiome function. Nucleic Acids Res 48:8883–8900.

Silva YP, Bernardi A, Frozza RL. 2020. The Role of Short-Chain Fatty Acids From Gut Microbiota in Gut-Brain Communication. Front Endocrinol 11:25.

Smith O, Dunshea G, Sinding M-HS, Fedorov S, Germonpre M, Bocherens H, Gilbert MTP. 2019. Ancient RNA from Late Pleistocene permafrost and historical canids shows tissue-specific transcriptome survival. PLoS Biol 17:e3000166.

Steinegger M, Söding J. 2017. MMseqs2 enables sensitive protein sequence searching for the analysis of massive data sets. Nat Biotechnol 35:1026–1028.

Sudo N, Chida Y, Aiba Y, Sonoda J, Oyama N, Yu X-N, Kubo C, Koga Y. 2004. Postnatal microbial colonization programs the hypothalamic-pituitary-adrenal system for stress response in mice: Commensal microbiota and stress response. J Physiol 558:263–275.

Suzek BE, Wang Y, Huang H, McGarvey PB, Wu CH, UniProt Consortium. 2015. UniRef clusters: a comprehensive and scalable alternative for improving sequence similarity searches. Bioinformatics 31:926–932.

Suzuki TA, Ley RE. 2020. The role of the microbiota in human genetic adaptation. Science 370. doi:10.1126/science.aaz6827

Teullet S, Tilak M-K, Magdeleine A, Schaub R, Weyer NM, Panaino W, Fuller A, Loughry WJ, Avenant NL, de Thoisy B, Borrel G, Delsuc F. 2023. Metagenomics uncovers dietary adaptations for chitin digestion in the gut microbiota of convergent myrmecophagous mammals. mSystems 8:e0038823.

Tikhonov G, Opedal ØH, Abrego N, Lehikoinen A, de Jonge MMJ, Oksanen J, Ovaskainen O. 2020. Joint species distribution modelling with the r-package Hmsc. Methods Ecol Evol 11:442–447.

Tovar A, Tews JK, Torres N, Harper AE. 1988. Some characteristics of threonine transport across the blood-brain barrier of the rat. J Neurochem 51:1285–1293.

Uritskiy GV, DiRuggiero J, Taylor J. 2018. MetaWRAP-a flexible pipeline for genome-resolved metagenomic data analysis. Microbiome 6:158.

Voolstra CR, Ziegler M. 2020. Adapting with microbial help: Microbiome flexibility facilitates rapid responses to environmental change. Bioessays 42:e2000004.

Warda AK, Rea K, Fitzgerald P, Hueston C, Gonzalez-Tortuero E, Dinan TG, Hill C. 2019. Heat-killed lactobacilli alter both microbiota composition and behaviour. Behav Brain Res 362:213–223.

Watanabe N, Mikami K, Hata T, Kimoto K, Nishino R, Akama F, Yamamoto K, Sudo N, Koga Y, Matsumoto H. 2021. Effect of gut microbiota early in life on aggressive behavior in mice. Neurosci Res. doi:10.1016/j.neures.2021.01.005

Wu Y-W, Simmons BA, Singer SW. 2016. MaxBin 2.0: an automated binning algorithm to recover genomes from multiple metagenomic datasets. Bioinformatics 32:605–607.

Yao Y, Han W. 2022. Proline Metabolism in Neurological and Psychiatric Disorders. Mol Cells 45:781–788.

Yarlagadda K, Zachwieja AJ, de Flamingh A, Phungviwatnikul T, Rivera-Colón AG, Roseman C, Shackelford L, Swanson KS, Malhi RS. 2022. Geographically diverse canid sampling provides novel insights into pre-industrial microbiomes. Proc Biol Sci 289:20220052.

Zhang X, Yoshihara K, Miyata N, Hata T, Altaisaikhan A, Takakura S, Asano Y, Izuno S, Sudo N. 2022. Dietary tryptophan, tyrosine, and phenylalanine depletion induce reduced food intake and behavioral alterations in mice. Physiol Behav 244:113653.

Zheng H, Xu P, Jiang Q, Xu Q, Zheng Y, Yan J, Ji H, Ning J, Zhang X, Li C, Zhang L, Li Y, Li X, Song W, Gao H. 2021. Depletion of acetate-producing bacteria from the gut microbiota facilitates cognitive impairment through the gut-brain neural mechanism in diabetic mice. Microbiome 9:145.

Zoelzer F, Burger AL, Dierkes PW. 2021. Unraveling differences in fecal microbiota stability in mammals: from high variable carnivores and consistently stable herbivores. Anim Microbiome 3:77.

