## Supplementary figures for "Functional insights into the effect of feralisation on the gut microbiota of cats worldwide"

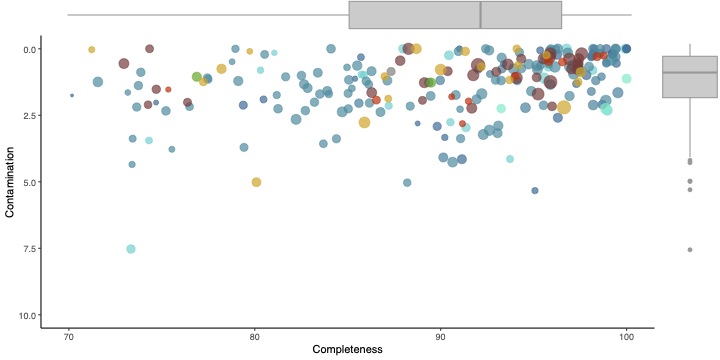

**Figure S1**. CheckM completeness and contamination of reconstructed metagenome-assembled genomes.

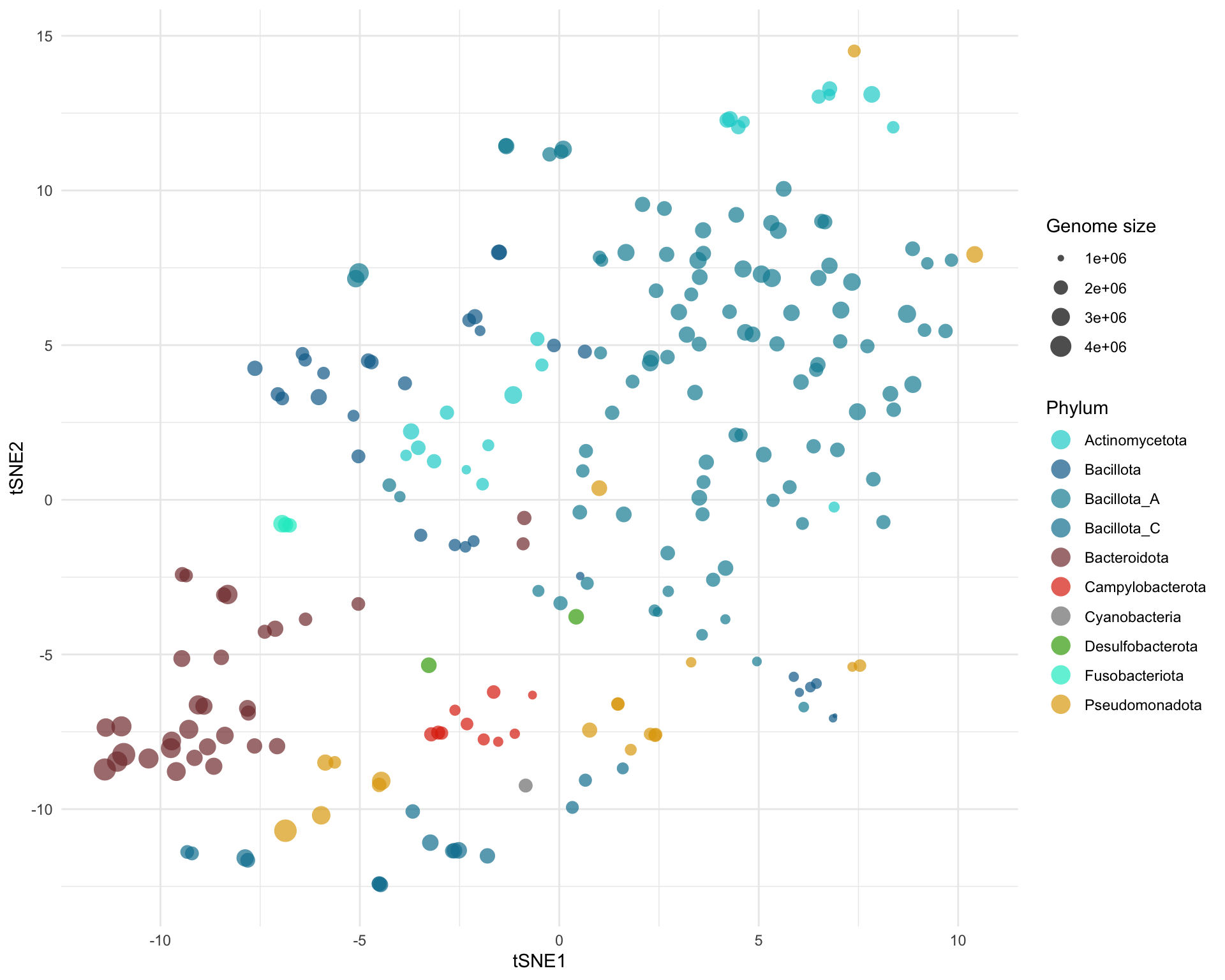
**Figure S2**. tSNE ordination of the metagenome-assembled genomes based on 170 genome-inferred functional traits (GIFTs).

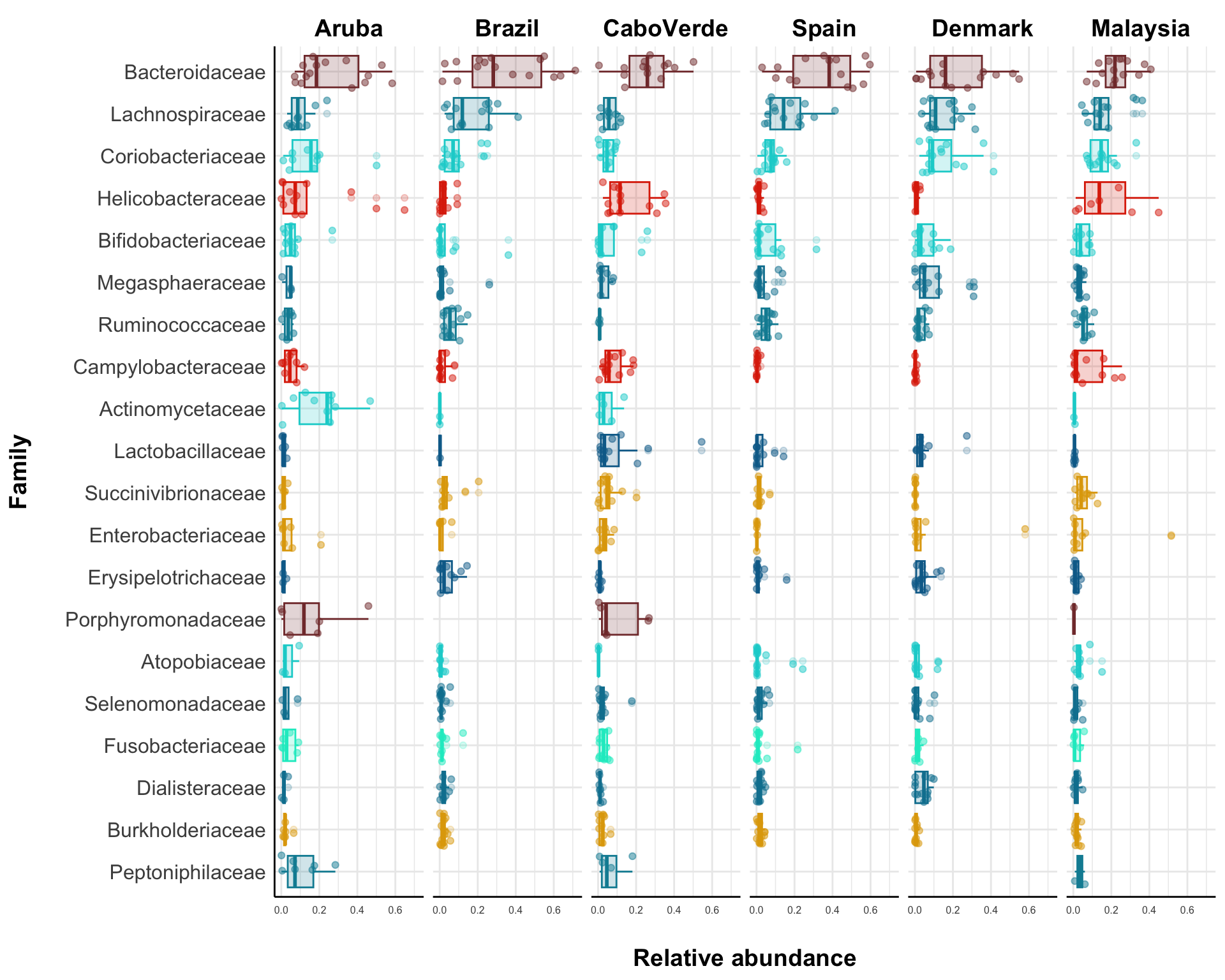
**Figure S3**. Average relative abundances of bacterial families across the six studied geographical regions.

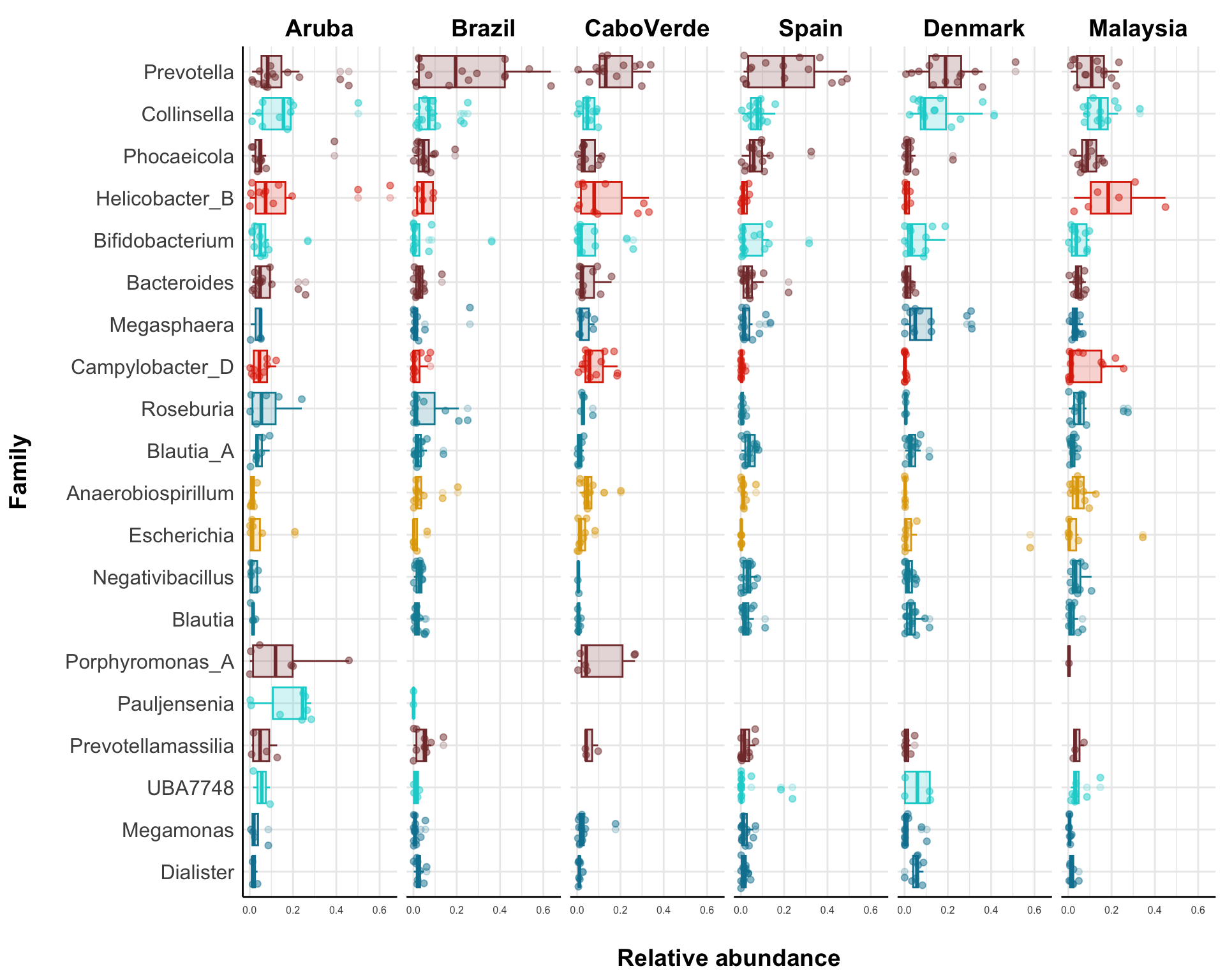
**Figure S4**. Average relative abundances of bacterial genera across the six studied geographical regions.

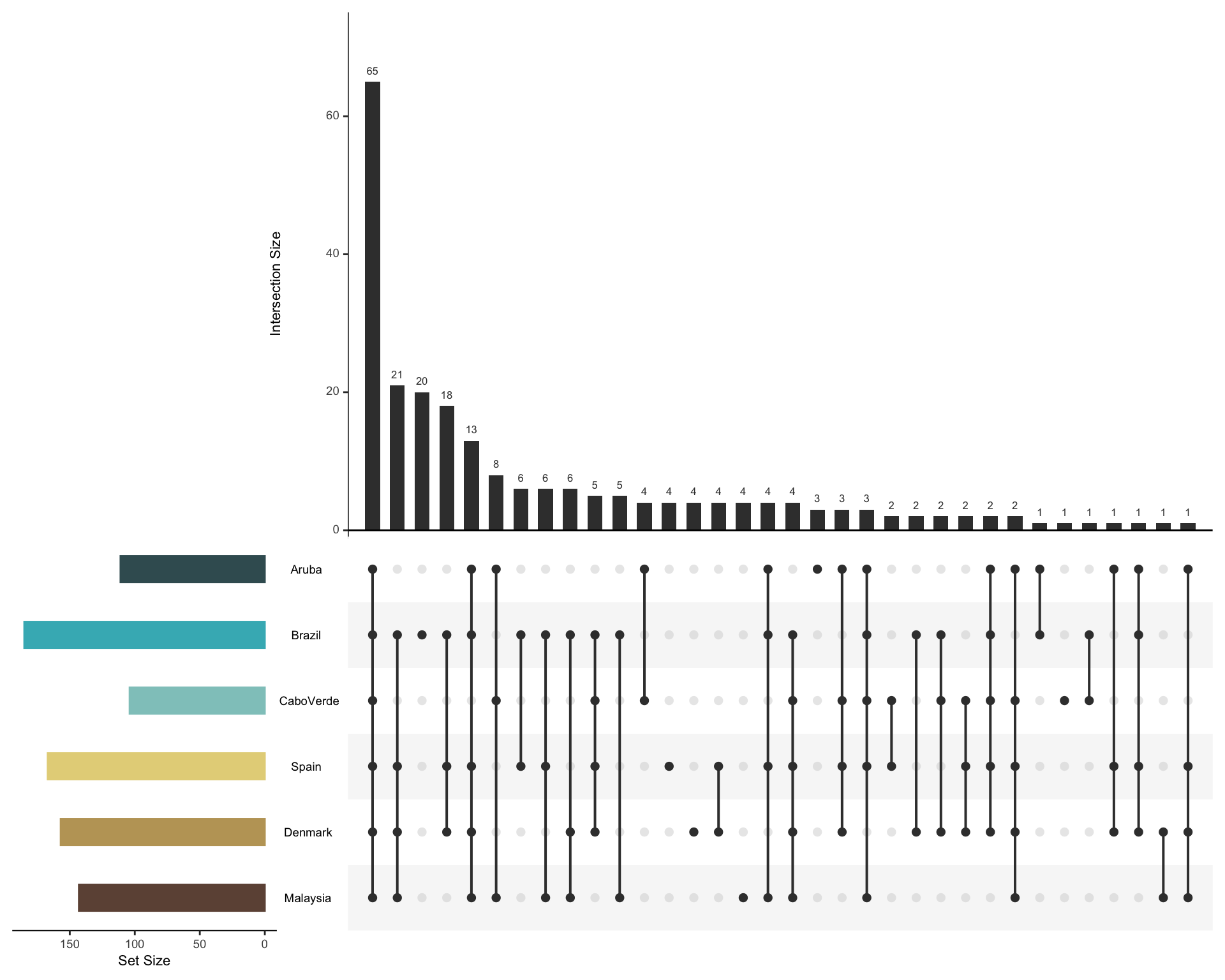

**Figure S5**. Prevalence of metagenome-assembled genomes across studied geographic regions.

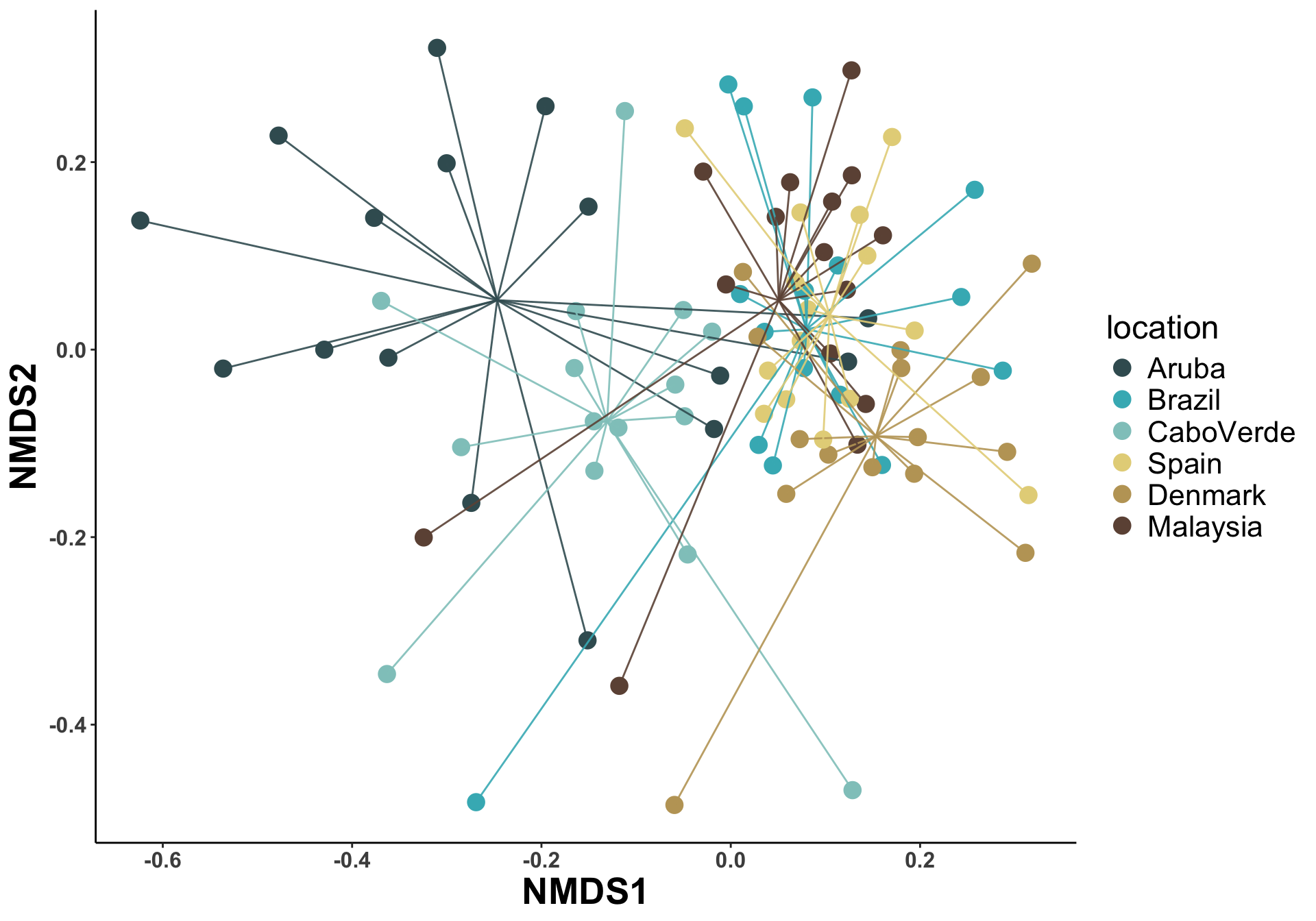

**Figure S6**. Compositional differences between samples measured based on beta diversities of neutral Hill numbers.
